# Gamma radiation-induced molecular toxicity and effects on pluripotent stem cells of the radiosensitive conifer Norway spruce (*Picea abies*)

**DOI:** 10.1101/2025.06.02.653021

**Authors:** Payel Bhattacharjee, YeonKyeong Lee, Marcos Viejo, Gareth B Gillard, Simen Rød Sandve, Torgeir R. Hvidsten, Brit Salbu, Dag A Brede, Jorunn E Olsen

## Abstract

Conifers are among the most radiosensitive plant species. Elevated, sublethal levels of ionising radiation result in reduced apical dominance in conifers, indicating a negative effect on shoot apical meristems (SAMs). The SAMs, harbouring the pluripotent stem cells, generate all the cells of the shoot, enabling growth and reproduction. However, knowledge on the effects of ionising radiation on such stem cells is scarce, but important for risk assessment and radioprotection of plants in contaminated ecosystems. Here, we assessed the sensitivity of *in vitro*-grown stem cells of Norway spruce to 144-h of gamma irradiation at 1-100 mGy h^-1^, using such cells as a model for molecular toxicity of gamma radiation in conifers. Although there were no visible effects of the gamma irradiation on cell proliferation and subsequent embryo formation, dose rate-dependent DNA damage was observed at ≥10 mGy h^-1^, and comprehensive organelle damage at all dose rates. Massive dose rate-dependent transcriptome changes occurred, with downregulation of a range of genes related to cell division, DNA repair and protein folding but upregulation of stress-related hormonal pathways and several antioxidant-related genes. The upregulation of such genes, survival and continued proliferation of at least a subset of cells and the post-irradiation normalisation of expression of DNA repair and protein-folding genes together with somatic embryo formation suggest that stem cells are able to recover from gamma-irradiation-induced stress. Collectively, regardless of cellular abnormalities after gamma irradiation, and huge transcriptomic shifts towards stress management pathways, the pluripotent stem cell cultures were able to retain their stemness.

**Highlights:**

- Norway spruce stem cells retained stemness after gamma irradiation (1-100 mGy h^-1^).
- DNA damage was observed at ≥ 10 mGy h^-1^ and organelle damage at all dose rates.
- Huge dose rate-dependent transcriptomic changes occurred after 144-h irradiation.
- Downregulated key DNA repair-genes recovered post-irradiation.
- Induction of stress-management and antioxidant genes may aid stem cell maintenance.

## Introduction

The extremely high ionising radiation doses from radioactive particles released from the Chernobyl nuclear power plant (ChNPP) accident on 26^th^ April 1986 resulted in massive mortality of the conifer tree species Scots pine (*Pinus sylvestris* L.) in a nearby 4.5 km^2^ forest area (Kashparova et al., 2020). In addition, elevated, sublethal radiation in the Chernobyl surroundings and in the Fukushima nuclear power plant accident area (2011, Japan) resulted in needle chlorosis, necrosis and reduced apical dominance leading to a bushy appearance in conifer trees such as Scots pine (*Pinus sylvestris* L.), Norway spruce (*Picea abies* L. (H). Karst), Japanese red pine (*Pinus densiflora* Sieb. et Zucc.) and Japanese fir (*Abies firma* Sieb. et Zucc.), (Kashparova et al. 2020; Geraskin et al. 2008; Watanabe et al. 2015; Yoschenko et al. 2016). The reduced apical dominance implies a negative effect on the shoot apical meristems (SAMs). After more than three decades, in 2018, Scots pine trees in sites with elevated radiation near the ChNPP still showed reduced apical dominance and increased DNA damage in shoot tips compared to trees grown in sites with background radiation (Nybakken et al. 2023).

The SAM and the root apical meristem (RAM), which are laid down during the embryogenesis, harbour the pluripotent stem cells that continuously generate cells forming vegetative organs under permissive environmental conditions and retain their stemness throughout the life of the plant (Umeda et al. 2021). The stem cells of the SAM also generate the reproductive meristems that are the source of the female and male gametophytes, which eventually contribute to the formation of embryos after the fertilisation. The high mutation rates in Scots pine progeny growing under chronic ionising radiation in the Chernobyl area (Caplin and Willey 2018), support adverse effects on the pluripotent cells of the SAMs, the reproductive meristems and embryo development. However, detailed information about the radiosensitivity of plant stem cells and effect on their stemness is limited.

Conifers are among the most radiosensitive plant species, while grasses and *Arabidopsis thaliana* are highly tolerant (Barescut et al. 2011; Sparrow and Miksche 1961; Blagojevic et al. 2019a and b; Bhattacharjee et al. 2025). Under laboratory conditions, gamma irradiation experiments employing up to 540 mGy h^-1^ from a ^60^Co source for 144 h, resulted in growth inhibition in Norway spruce and Scots pine seedlings at ≥ 20-40 mGy h^-1^, more disorganised SAMs with increasing dose rate, and mortality at the highest dose rates (Blagojevicet al. 2019a). By contrast, only a slight delay in lateral root formation and flower bud appearance at dose rates ≥ 290-400 mGy h^-1^ was observed in *A. thaliana*, with no visible negative effects on the SAMs, and no mortality. Moreover, significant DNA damage was observed in all three species exposed to dose rates from 1 mGy h^-1^, but with more DNA damage in the conifers than in *A. thaliana* (Blagojevic et al. 2019a). Ionising radiation can result in damage to vital cellular macromolecules such as nucleic acids, lipids and proteins through direct ionisation and the formation of reactive oxygen species (ROS) (You and Chan 2015). Several studies, particularly in *A. thaliana*, have indicated the activation of DNA damage repair (DDR) and antioxidants protecting towards oxidative stress in plants during exposure to ionising radiation (Kim et al. 2019). The DDR encompasses a complex cascade of events including checkpoint activation, cell cycle arrest or endoreduplication, activation of different DDR pathways and programmed cell death (PCD). A recent comparative study suggested more efficient transcriptional mobilisation of DDR and antioxidants in gamma-irradiated seedlings of *A. thaliana* than Norway spruce (Bhattacharjee et al. 2025). However, despite that stem cells in the SAMs and RAMs are crucial for vegetative and reproductive plant development in natural and man-made ecosystems, the effect of ionising radiation on repair and protection systems in stem cells has previously not been investigated. Additionally, stressful conditions typically lead to stiffening of the plant cell wall through interaction between stress-induced ROS formation, cell wall integrity sensing and hormone signalling pathways (Novaković et al. 2018; Tenhaken 2014). However, the knowledge on how ionising radiation affects the plant cell wall composition is limited.

A powerful model system for pluripotent stem cells of conifers can be induced *in vitro* from e.g., excised zygotic embryos or primordial shoot explants (Varis, Klimaszewska, and Aronen 2018). Such stem cells can retain their stemness for a very long time and generate somatic embryos upon induction treatment, rendering it a valuable tool to study embryogenesis in plants with long generation times such as conifers, e.g., about 25 years in Norway spruce.

In this study, *in vitro*-grown pluripotent stem cells of Norway spruce were employed to analyse radiotoxicity in ecologically and economically important conifers, and specifically to test the hypothesis that low to moderate levels of ionising radiation affects the viability and stemness of stem cells in conifers. Given the importance of the stem cell-harbouring apical meristems for growth, reproduction and survival of plants, such information is important for predicting responses of ionising radiation in ecosystems, and accordingly for risk assessments and radioprotection. Here, stem cells of Norway spruce were gamma-irradiated and the damage and repair mechanisms as well as effects associated with the molecular cell physiology were investigated.

## Materials and Methods

### Plant materials and pre-growing conditions

*In vitro* stem cell cultures of Norway spruce (*Picea abies* (L.) H. Karst.) were generated following the protocol by (Kvaalen and Johnsen 2008). This included the following steps: A single zygotic seed originating from controlled crosses of the parents #26509 and #2707 (Johnsen et al. 2005) was used to initiate cultures of genetically identical cells (pluripotent stem cells) denoted as clone B10V. Cell proliferation media (PM) with the following composition was used: AL medium (formulated as a basal medium for *in vitro* culture of *Abies lasiocarpa*) (Kvaalen et al. 2005) supplemented with inositol (10 %, w/v), a revised vitamin mixture previously used for *in vitro* culture of Douglas-fir cotyledons (Gupta and Durzan 1985), 2,4 dichlorophenoxy acetic acid (2,4-D; 10 μM), benzyl aminopurine (BA; 5 μM), sucrose (1 %, w/v), and Phytagel (P-8169; Sigma-Aldrich, Steinheim, Germany; 0.3 % w/v) (Yakovlev et al. 2014). The proliferating cells were kept as cell aggregates on PM in Petri dishes of 9 cm diameter at 23°C in darkness. The cell aggregates (10 per dish) were divided and transferred to fresh media every two weeks until an adequate number of aggregates was obtained for the gamma irradiation experiments.

### Gamma irradiation of stem cells and growing conditions during the irradiation

The FIGARO low dose gamma irradiation facility (^60^Co; 1173.2 and 1332.5 keV γ-rays) at the Norwegian University of Life Sciences (Lind et al. 2018) was used to expose aggregates of proliferating stem cells on PM to gamma radiation at 1, 10, 20, 40 and 100 mGy h^-1^ for 144 h (6 days) in the dark at 23°C ±1°C. To obtain these dose rates, the Petri dishes containing densely spaced, not overlapping cell aggregates, were placed at different distances from the ^60^Co source and rotated 180° halfway through the experiment to ensure that all cells received as even gamma radiation as possible. The dose rates to water and dose rate intervals (Table 1) were calculated from the air kerma rates according to (Hansen et al. 2019) as previously described (Blagojevic et al. 2019a and b). Two Petri dishes containing 10-15 cell aggregates each per dose rate were exposed in each of two repeated experiments, with unexposed control samples kept in the same room outside the radiation sector, behind lead shields. Thus, per dose rate, altogether 40 or more cell aggregates were exposed.

**Table 1.** The gamma radiation treatments applied in the experiments with exposure of stem cells of Norway spruce for 144 h using a ^60^Co source.

| Dose rate (mGy h<sup>-1</sup>) | Dose rate interval (mGy h<sup>-1</sup>) |  | Total dose (mGy) |
| --- | --- | --- | --- |
|  | Min | Max |  |
| 0 | 0.002 | 0.006 | 0,576 |
| 1 | 1.0 | 1.1 | 144 |
| 10 | 10.5 | 10.6 | 1440 |
| 20 | 18.7 | 21.0 | 2880 |
| 40 | 33.3 | 43.9 | 5760 |
| 100 | 83.8 | 114.8 | 14400 |

### Post-irradiation growing conditions

To evaluate the post-irradiation cell proliferation, gamma irradiated cell aggregates were maintained in PM at 23°C in darkness for 7 weeks with change of media and division of the cell aggregates every two weeks. Proliferation stage 1 (P1) and 2 (P2) denote the stages 5 and 7 weeks on PM after the gamma irradiation, respectively. Thereafter, to study the after-effects of the gamma irradiation on subsequent embryogenesis, cell aggregates were transferred to maturation media (MM), consisting of PM without the growth regulators 2,4-D and BA, but supplemented with 20 μM abscisic acid (ABA; A8451, Sigma-Aldrich). The carbon source was changed from sucrose to maltose and the medium was supplemented with polyethylene glycol (PEG4000; Fluka, Switzerland) (5 %, 10 % and 13 % w/v in MM in stage 1, 2 and 3, respectively) (Yakovlev et al. 2014). The cell aggregates were transferred to fresh MM every two weeks. Maturation stage 1 and 2 denote the stages 11 and 15 weeks after the gamma irradiation, respectively, corresponding to 4 and 8 weeks after the transfer to MM. Maturation stage 3-1, 3-2 and 3-3 denote the stages 17, 20 and 24 weeks after the gamma exposure, corresponding to 10, 13 and 17 weeks on MM.

### COMET assay for analysis of DNA damage

To quantify the DNA damage after the gamma irradiation, the COMET assay was used following the protocol of (T. Gichner 2003) with some modifications as described by (Blagojevic et al. 2019a). To avoid light-induced DNA damage, the assay was performed under inactinic red-light conditions. Briefly, the protocol included isolation of cell nuclei from approximately 200 mg cells per treatment and electrophoresis in agarose gels under alkaline pH, followed by quantification of the DNA in the COMET tail (elongated cell nuclei due to DNA damage) using fluorescence microscopy. Per gamma dose rate, three repeated samples with 65-70 nuclei per sample were investigated individually for DNA damage using three gels per sample. The median value for each sample was calculated, followed by calculation of the average value for the three repeated samples, according to (Koppen et al. 2017).

### Studies of cell and organelle integrity by light microscopy and transmission electron microscopy

To investigate effects of gamma irradiation on the cell and organelle integrity in the stem cell aggregates at the end of the gamma irradiation and selected post-irradiation time points during the proliferation (5 and 7 weeks) and maturation (11, 15, 17, 20 and 24 weeks) stages, samples were harvested for histological and cytological studies. For each dose rate and developmental stage, samples from 3 different cell aggregates were harvested, fixed and embedded in LR White resin (London Resin Company, England) according to Lee et al. (2017). For light microscopy studies, 1 µm thick sections were made from embedded samples using an Ultracut Leica EM UC6 microtome (Leica, Mannheim, Germany), stained with toluidine blue O to visualise the cells and inspected using a light microscope with bright field optics (DM6B, Leica). For cytological observations, LR white-embedded samples were sectioned into 70 nm ultrathin sections, using an Ultracut microtome (Leica EMUC6) and stained with uranyl acetate and lead citrate before examining by transmission electron microscopy (TEM) (FEI Morgagni 268, United States).

### Analysis of cell wall modifications using immunolabeling

To investigate cell wall modifications in response to the gamma irradiation, indirect immunolabeling of cell wall components were performed in a time course as described in (Lee et al. 2017) using 3 plants per dose rate. For this purpose, 1 µm-thick LR White sections were obtained as described above. The sections were incubated with primary rat monoclonal antibodies (Table S1), and then with secondary antibodies (anti-rat-IgG, whole molecule) linked to fluorescein isothiocyanate (FITC), following a standard protocol. Afterwards, the sections were washed with PBS buffer, mounted with Fluoro-Gel (EMS, US) and examined using a microscope equipped with epifluorescence (Leica DM6B).

### RNA extraction

At the end of the gamma irradiation (Time point 1 = T1) and 7 weeks thereafter during the post-irradiation cell proliferation period (proliferation stage 2 = P2), 8 and 4 repeated samples per dose rate, respectively, were harvested for RNA isolation by flash-freezing in liquid nitrogen and stored at -80°C until further analysis. For each dose rate, four of the T1-samples were used for RNA sequencing (RNA-seq), and four T1-samples and four P2-samples were used for RT-qPCR analysis of transcript levels of specific candidate genes.

Total RNA was extracted using the MasterPure Complete DNA and RNA Purification Kit (Epicenter, Madison, WI, USA), following the manufacturer’s protocol except that 1-2 % polyvinylpyrrolidone (PVP, mw 360 000, Sigma-Aldrich) was added to the extraction buffer and 1,4-dithiothreitol (DTT) was replaced with 3 µl beta-mercaptoethanol. The concentration of PVP was 1 % for the T1-samples and 2 % for the P2-stage samples due to higher phenolic content. The purified RNA was quantified using a NanoDrop ND-1000 Spectrophotometer (Thermo Scientific, Wilmington, DE, USA) and the quality was determined by an Agilent 2100 Bioanalyzer (Agilent Technologies, Palo Alto, CA, USA).

### RNA seq; differential expression analyses and functional term enrichment

The transcriptomes were sequenced on the Illumina HiSeq4000 platform at the Norwegian Sequencing Centre (University of Oslo, Oslo, Norway) using the Strand specific 20x Truseq RNA library preparation (paired end; with read length of 150 bp) on two lanes on the HiSeq 4000 instrument.

Analyses of the RNA-seq data were performed as described in Bhattacharjee et al. (2023). The transcript abundances were estimated using the Salmon package (Patro et al. 2017) comparing each dose rate with the Norway spruce genome (*Picea abies* 1.0 assembly, plantgenie.org (Nystedt et al. 2013)). Gene expression in the different gamma-irradiated samples was compared with the unexposed control samples using DESeq2. Genes were classified as differentially expressed genes (DEGs) if the False Discovery Rate (FDR) (Benjamini & Hochberg correction) adjusted p-values were < 0.05.

Gene ontology (GO) annotations of Norway spruce genes as well as spruce to *A. thaliana* orthologs by best sequence matches were downloaded from plantgenie.org (Sundell et al. 2015). Enrichment of GO terms or KEGG (Kyoto Encyclopedia of Genes and Genomes) pathways in sets of DEGs was performed using topGO (version 2.34.0) or kegga from the limma package (version 3.38.2), respectively. The *A. thaliana* orthologs were used to associate the Norway spruce genes with KEGG pathways.

### RT-qPCR analysis

For P2 and T1 samples, cDNA was synthesised from 1 µg of RNA in a 20 µl reaction using SuperScript™ VILO™ cDNA Synthesis Kit (Invitrogen, Carlsbad, CA, US) following the manufacturer’s protocol and no enzyme controls (-RT) were made for each sample. Eight genes belonging to the functional categories DNA repair and unfolded protein response (UPR), were selected (Table S2) for the RT-qPCR analysis, based on their significant regulation by gamma irradiation as revealed by the RNA-seq analyses. To design primers, gene sequences were retrieved from PlantGenIE (plantgenie.org) (Table S2). Primers were designed using the online tools Oligoanalyzer (eu.idtdna.com/calc/analyzer) and Primer Blast (ncbi.nlm.nih.gov/tools/primer-blast/) with default parameters. All amplifications were performed in a 7500 Fast Real-time PCR system (Applied Biosystems, Foster city, USA) using 20 μl of Platinum Quantitative PCR Supermix-UDG and SYBRGreen (Thermo Fisher, Carlsbad, CA, US).

To compare the relative transcript levels of the selected genes in the samples exposed to different dose rates of gamma radiation with the unexposed controls, the comparative C_T_ (Cycle threshold) method (known as ΔΔC_T_ method) was used (Livak and Schmittgen 2001). The expression of the target genes was normalised by the mean expression of the three reference genes, α*-TUBULIN*, *ACTIN*, *ELONGATION FACTOR1*α. Four biological replicates per dose rate with 4 technical replicates of each were used in the qPCR analyses.

### Statistical analyses of DNA damage and qPCR gene expression data

For statistical analyses of DNA damage and gene expression analysed by qPCR, one-way analyses of variance (ANOVA) in the general linear model mode, followed by Tukey’s post hoc test, was performed (pL≤L0.05) using the Minitab statistical software (Minitab 18, Minitab Inc, PA, USA). Prior to these analyses, tests for equal variance and normal distribution were performed using Levenès and Ryan-Joiner’s tests, respectively. Due to lack of normal distribution, the gene expression data were log10 transformed prior to the ANOVA. (Minitab 18).

## Results

### No visible effect of gamma irradiation on cell proliferation or subsequent embryogenesis

There were no signs of differential proliferation or any other visible changes in the stem cell aggregates for any of the gamma dose rates following the 144-h gamma irradiation (Fig. 1a). Also, during the post-irradiation proliferation and subsequent maturation stages after transfer to embryogenesis-induction media, there were no visible differences between the gamma-exposed and the control cells (Fig. 1a, Fig. S1). Somatic embryos developed post-irradiation regardless of gamma dose rate; equal numbers of embryos were obtained in the controls and at 100 mGy h^-1^ (Fig. S1, Table S3). However, normal embryos with 7-8 cotyledons (Alvarez et al. 2015) were observed in the unexposed control only (results not shown).

**Fig. 1.**
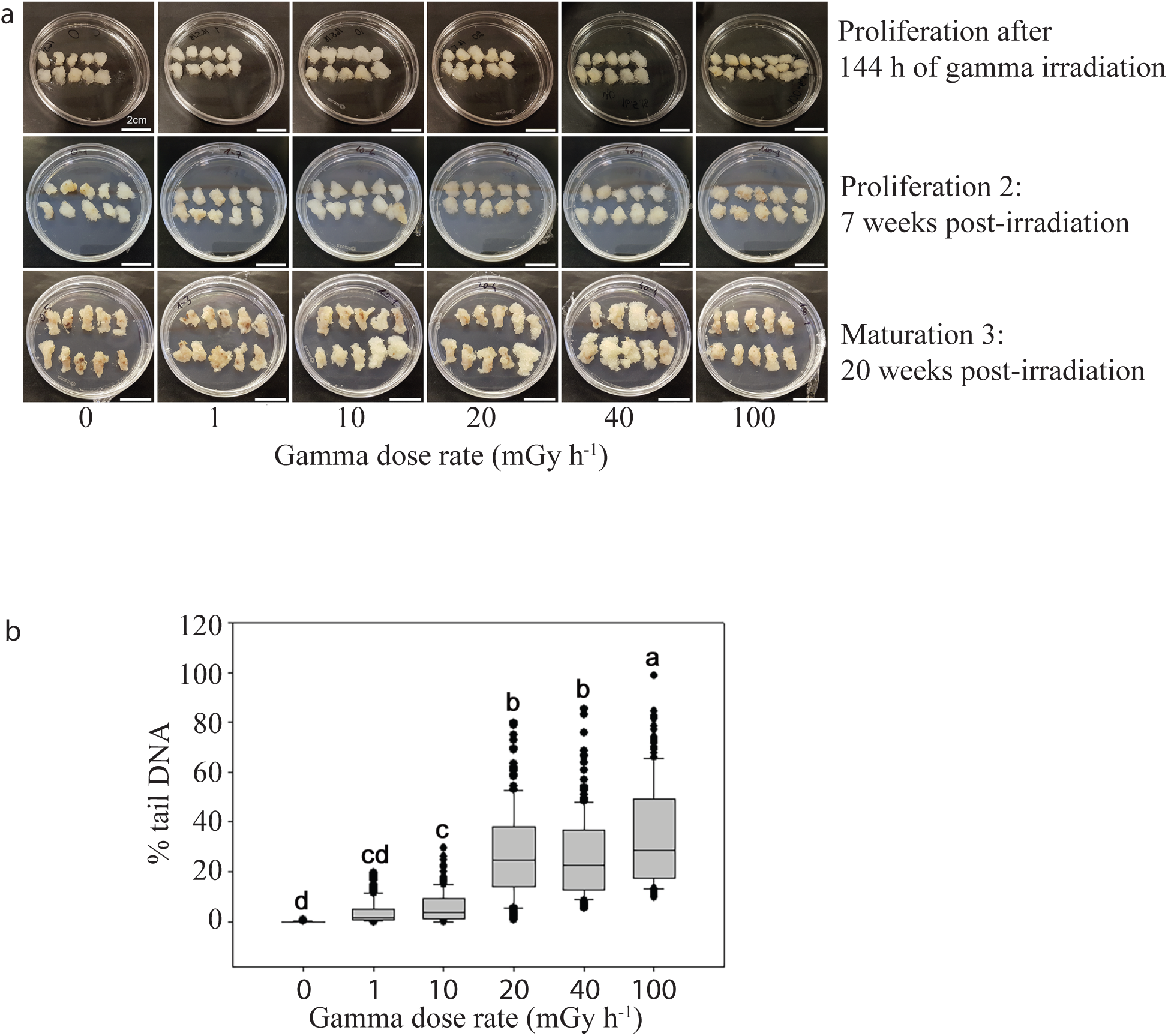
Effect of 144 h of gamma irradiation on clonal stem cells of Norway spruce. (a) Proliferating stem cell aggregates at the end of the exposure (upper panel) and during post-irradiation stages. Scale bars: 2 cm. (b) DNA damage after the gamma irradiation as measured by the COMET assay. The line in each box represents the median values of 3 biological replicates per dose rate with 3 technical replicates (gels) for each (n = 200 nuclei per gamma dose rate with 60-70 nuclei per biological replicate). Lower and upper boundaries = 25 and 75 percentiles, error bars = 10 and 90 percentiles with data points outside these shown as dots. Different letters indicate significant differences (p ≤ 0.05) based on analysis of variance followed by Tukey’s test.

### Dose rate-dependent DNA damage in gamma-irradiated cells

COMET analyses revealed significant DNA damage at ≥ 10 mGy h^-1^ (Fig. 1b). In response to 10, 20, 40 and 100 mGy h^-1^ the cells showed 5.8, 27.2, 26.3 and 34.9% tail DNA (Fig. 1b). At 100 mGy h^-1^, the DNA damage level was about 330 times higher than in the control (0.1 % tail DNA) and about 10 times higher than at 1 mGy h^-1^ (3.8 %).

### Gamma radiation-induced cell and organelle damage at all dose rates

Light microscopy (Fig. 2a) and TEM (Fig. 2b) analyses revealed cellular abnormalities such as disintegrating nuclei and nuclear membranes, disarrayed mitochondrial cristae and accumulation of amorphous materials in the cells at all gamma radiation dose rates at the end of the irradiation as well as during the post-irradiation stages. Generally, as expected, the cells gradually developed a more compact structure with smaller vacuoles, although exceptions with larger vacuoles were also observed, particularly for gamma-irradiated cells with different cellular abnormalities (Fig. 2a).

**Fig. 2.**
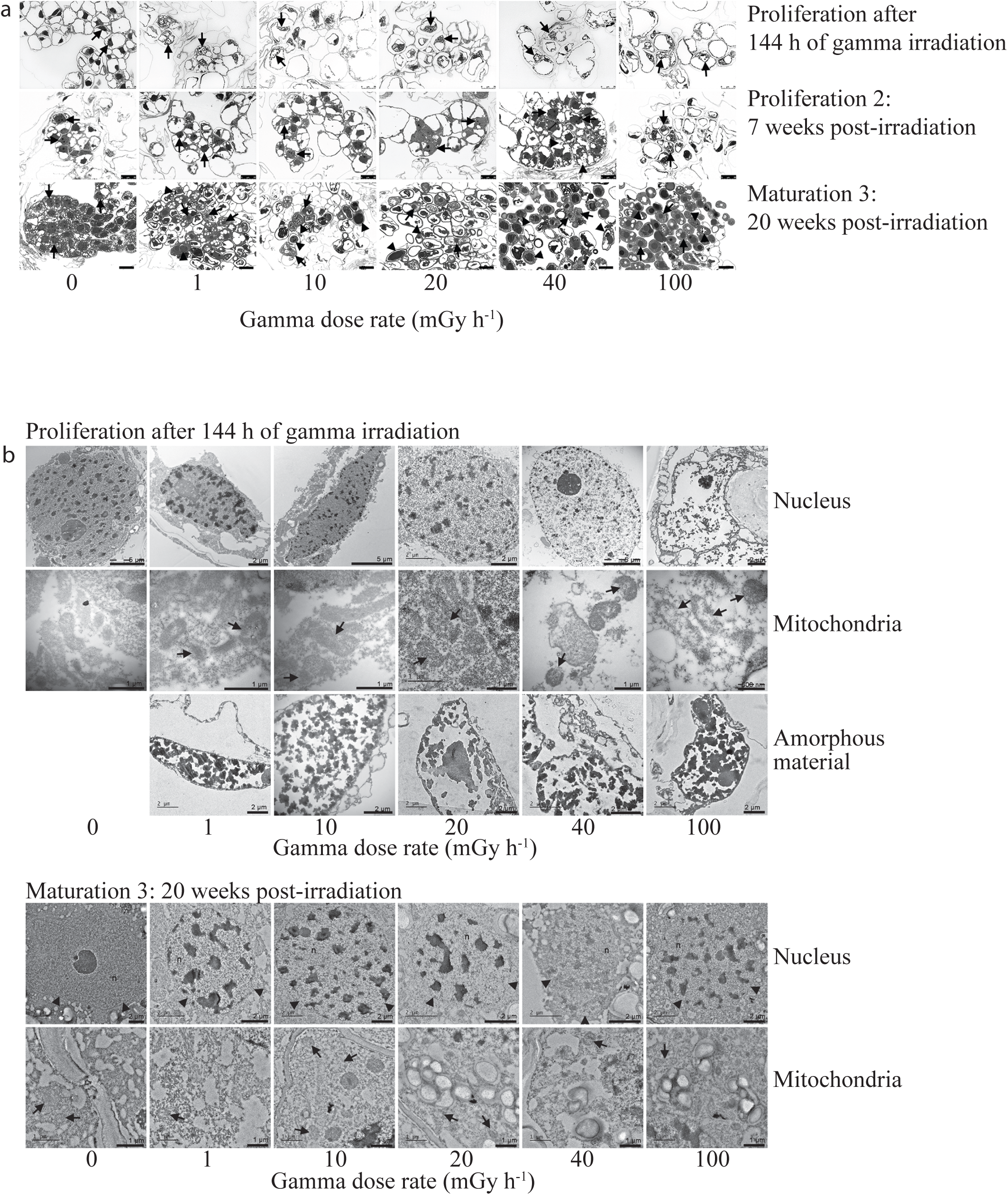
Cellular damage in clonal stem cells of Norway spruce after 144 h of gamma irradiation. (a) Light micrographs at the end of the exposure (upper panel) and during post-irradiation proliferation and maturation stages (middle and lower panel). Arrows indicate cell nuclei and arrowheads indicate amorphous materials. Scale bars: 25 µm. (b) Transmission electron micrographs of stem cells of Norway spruce at the end of the gamma irradiation (upper three panels) and during a post-irradiation maturation state. Arrows indicate normal control cells and cellular abnormalities in gamma-exposed cells.

### Cell wall modifications in response to gamma radiation

At the end of the gamma-irradiation no immunofluorescence signal was detected for the methyl-esterified homogalacturonan (HG) pectin using the LM20 antibody, and the control cells exhibited a weak signal only (Fig. S2a). However, during the post-irradiation culture period both the unexposed control and gamma-irradiated cells showed similar, stronger signals. The abundance of un-methyl-esterified HG detected by the LM19 antibody did not differ between the 144 h-gamma-irradiated and control cells, but at the post-irradiation stages (from 7 weeks), the immunofluorescence signal was generally stronger in the gamma-irradiated cells (Fig. S2b). The signal for the hemicellulose xyloglucan detected by the LM15 antibody appeared similar at all gamma radiation dose-rates both at the end of the gamma irradiation and post-irradiation but there were no or very low signals only in the unexposed control cells (Fig. S2c). The arabinogalactan protein detected by the JIM13 antibody was observed in gamma-irradiated cells during late post-irradiation maturation stages only but not in control cells at any stage (Fig. S2d).

### Substantially altered transcriptomes in response to gamma-irradiation

RNA-seq analysis revealed that the number of differentially expressed genes (DEGs) between the gamma-irradiated cells (144 h) and the unexposed controls increased with increasing dose rate (Fig. 3a). Amongst 66069 predicted genes in Norway spruce, 17047 (25 %) were DEGs (FDR adjusted P-values < 0.05) (Fig 3b). Although there was a substantial number of unique DEGs for each dose rate, some overlap between the dose rates was observed (Fig 3c). Moreover, 15365 genes showed significant difference in expression at 100 mGy h^-1^ compared to the unexposed controls. GO (Fig. S3) and KEGG pathway (Fig. S4) enrichment analyses showed that cell cycle/cell division, DNA repair mechanisms, DNA methylation-dependent chromatin silencing-related genes, and selected pathways involved in development were enriched for DEGs. GO categories related to initiation and regulation of DNA replication, G2/M transition and cell proliferation were enriched for downregulated DEGs (Fig. S3), indicating that regulation of cell cycle and replication must be considerably hindered due to radiation stress. Additionally, KEGG enrichment analyses found DNA repair pathways such as base excision repair, nucleotide excision repair and mismatch repair among the most enriched pathways for downregulated DEGs (Fig. S4). A wide range of genes encoding transcription factors, hormonal pathways, protein folding, protein degradation and transport and secondary cell wall biogenesis also demonstrated modified transcriptional patterns upon the radiation stress (Table 2-7, additional information and details in Table S4-S11).

**Fig. 3.**
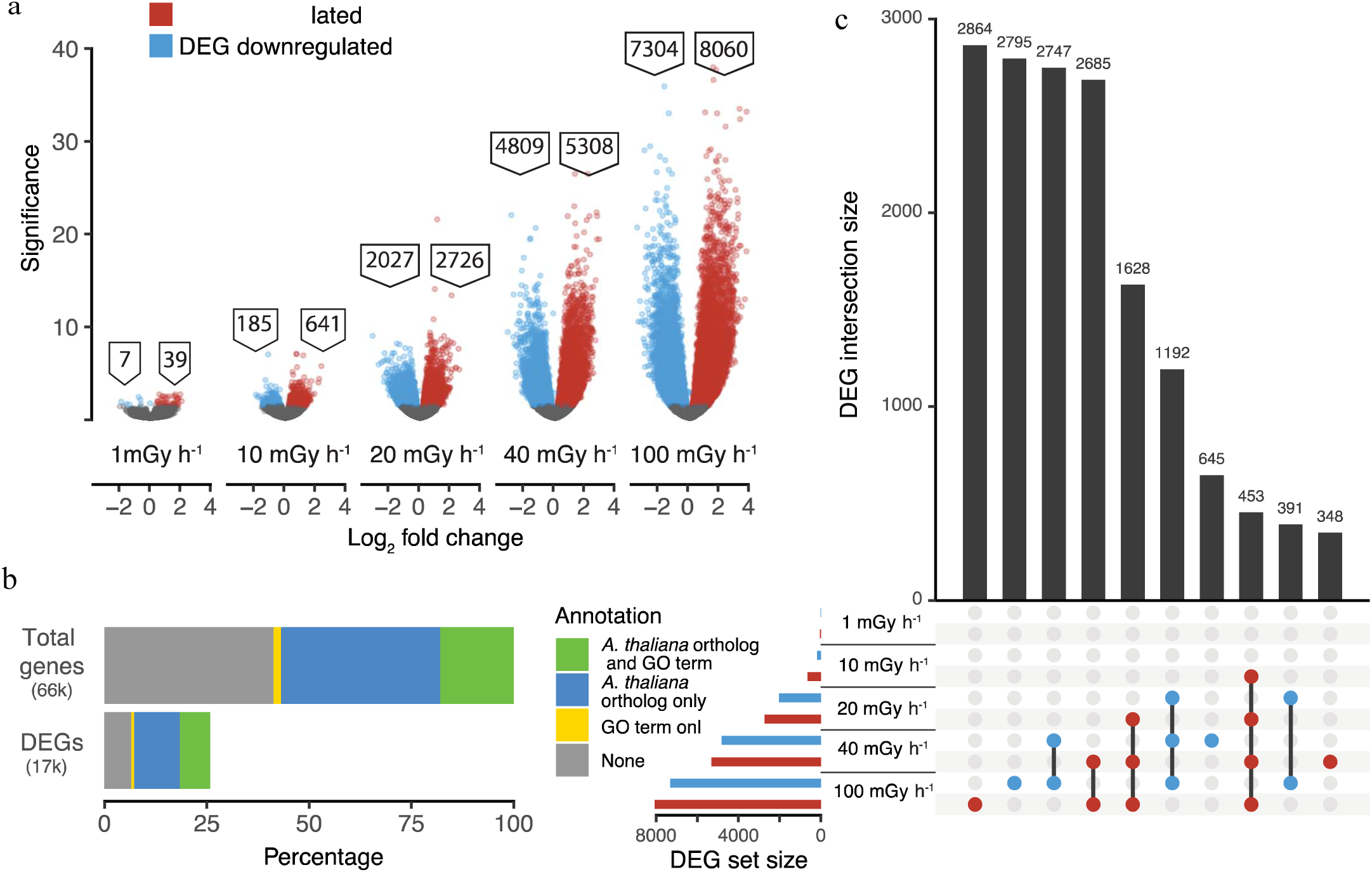
Effect of different gamma dose rates for 144 h on gene expression in clonal stem cells of Norway spruce. For each dose rate four repeated samples were analysed in duplicate by RNA sequencing. (a) Number of differentially expressed genes (DEGs; log_2_ fold change in DEGs, adjusted p-value < 0.05) with up- (red) or down- (blue) regulated genes relative to the unexposed control (0 mGy h^-1^). (b) Upset plot showing the numbers of up- (red) or down- (blue) regulated DEGs and overlap between the different gamma dose rates. (c) Percentages of the total number of genes and DEGs (25% of the total) with available annotation data (Gene Ontology (GO) terms or an *Arabidopsis thaliana* gene ortholog).

**Table 2.**
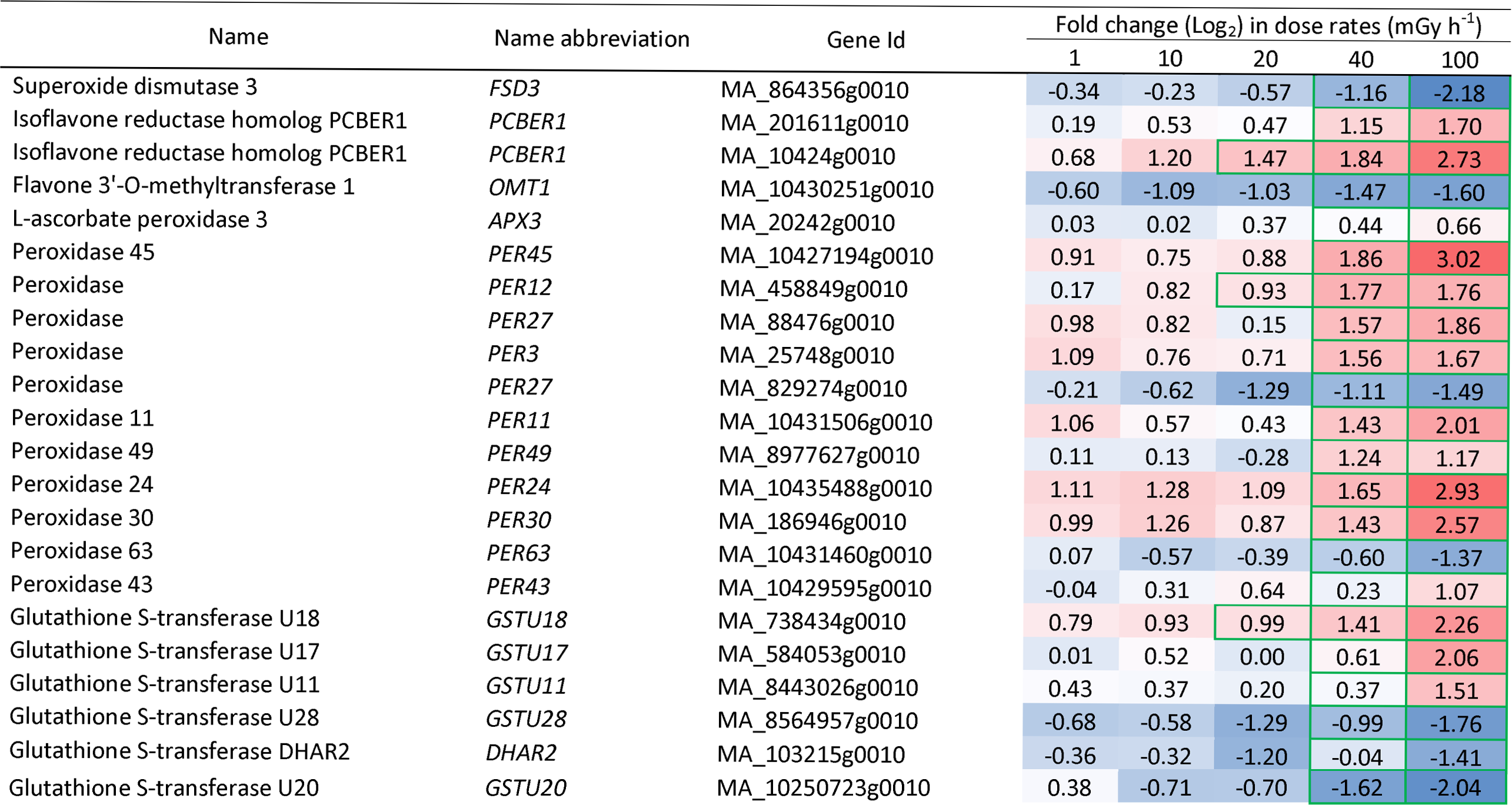
Differentially expressed antioxidant-related genes in clonal stem cells of Norway spruce after exposure to 144 h of gamma radiation at different dose rates. Red and blue highlights denote upregulation and downregulation, respectively, compared to unexposed control (0 mGy h^-1^). Green borders denote significant fold changes (p□≤□0.05).

### Gamma radiation-induced changes in expression of genes related to antioxidants, DDR, cell cycle regulation and epigenetics

A wide range of antioxidant biosynthesis and signalling genes were significantly regulated in cells exposed to dose rates ≥ 40 mGy h^-1^ (Table 2, Table S4). Among others, a range of *PEROXIDASEs* (*PER*s)*, GLUTATHIONE-S-TRANSFERASEs* (*GSTU*s) and *PHENYLCOUMARAN BENZYLIC ETHER REDUCTASE 1* (*PCBER1*), showed upregulation mostly at 40-100 mGy h^-1^. *SUPEROXIDE DISMUTASEs* (*SOD*s), on the other hand, were downregulated at ≥ 40 mGy h^-1^.

More than 250 DDR-related genes were significantly regulated after the gamma irradiation with approximately 95 % showing downregulation, particularly at 100 mGy h^-1^ (Table 3, Table. S5). *SUPRESSOR OF GAMMA RESPONSE 1* (*SOG1*) that encodes a master regulator of DNA damage-induced cell cycle arrest in plants, was downregulated at all dose rates and decreased with increasing dose rate. *RAD4*, *RAD5A* and *RAD50* were downregulated at 100 mGy h^-1^ (Table 3). The DNA repair protein *X-REPAIR CROSS COMPLEMENTING 3* (*XRCC3*) gene was one of the few DEGs that showed upregulation.

**Table 3.**
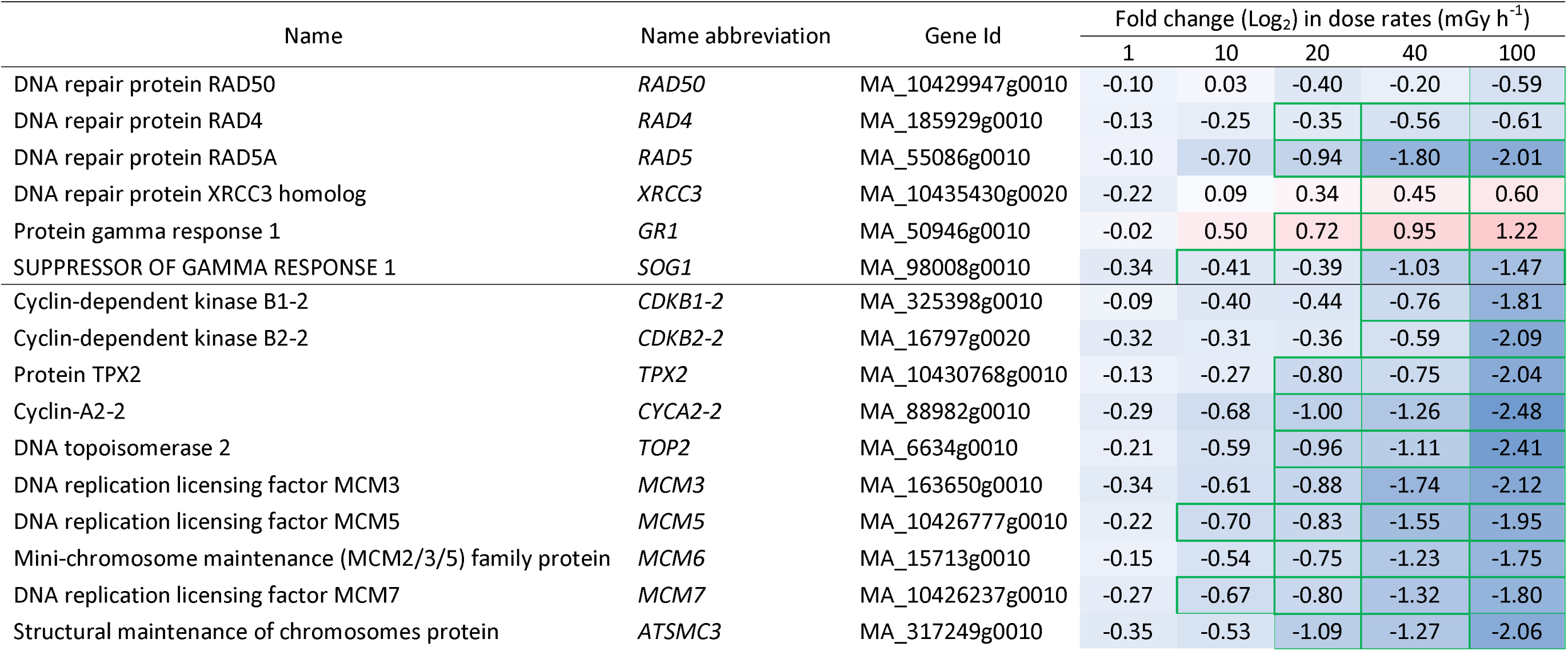
Differentially expressed DNA repair and cell cycle related genes in clonal stem cells of Norway spruce after exposure to 144 h of gamma radiation at different dose rates. Red and blue highlights denote upregulation and downregulation, respectively, compared to unexposed control (0 mGy h^-1^). Green borders denote significant fold changes (p□≤□0.05).

Significant downregulation of cell-cycle related genes was observed at dose rates ≥ 40 mGy h^-1^ (Table 3, Table S6). The pro-spindle assembly factor *TARGETING PROTEIN FOR XKLP2* (*TPX2*), *CYCLIN A2* (*CYC2A2*) and the cyclin-dependent kinase genes, *CDKB1* and *CDKB2*, were downregulated at ≥ 40 mGy h^-1^ (Table 3, Table S6). *DNA TOPOISOMERASE 2* (*TOP2*) and multiple genes encoding components of the mini chromosome maintenance protein complex (MCM; a DNA helicase), and *STRUCTURAL MAINTENANCE OF CHROMOSOME 3* (*SMC3*) showed dose responsive downregulation at ≥ 20 mGy h^-1^ (Table 3).

GO terms related to DNA modification were significantly enriched, and nearly 200 genes with predicted function related to epigenetic pathways or chromatin modifications were significantly regulated in a dose rate-dependent manner (Fig. S3, Table 4, Table S7). Many genes encoding DNA methylases, helicases and histone chaperones showed ≥ 2-fold regulation. The ATP-dependent DNA helicase *DECREASE IN DNA METHYLATION 1* (*DDM1*) gene, *DMT1* (*DEFICIENT IN DNA METHYLATION 1*), *CHROMOMETHYLASE 2 AND 3* (*CMT2* and *CMT3*), were downregulated at ≥ 20 mGy h^-1^ and one of the histone chaperones *ANTI-SILENCING FACTOR 1B* (*ASF1B)* were downregulated at ≥ 40 mGy h^-1^ (Table 4, Table S7). A drastic change in expression of genes encoding the Histone 2A (H2A) and 2B (H2B) subunits were also observed with the H2 component genes *HTA5* and *HTB11* being downregulated at ≥ 10 mGy h^-1^ and at ≥ 20 mGy h^-1^, respectively (Table 4). In addition, the core histone-binding subunit gene *MULTICOPY SUPRESSOR OF IRA1* (*MSI1*) and the histone methyltransferase genes *SU(VAR)3-9 HOMOLOGUE 5 (SUVH5)* and *SUVR4* were downregulated from ≥ 20 mGy h^-1^ (Table 4).

**Table 4.** Differentially expressed Epigenetics or chromatin modification related genes in clonal stem cells of Norway spruce after exposure to 144 h of gamma radiation at different dose rates. Red and blue highlights denote upregulation and downregulation, respectively, compared to unexposed control (0 mGy h^-1^). Green borders denote significant fold changes (p□≤□0.05).

| Name | Name abbreviation | Gene Id | Fold change (Log<sub>2</sub>) in dose rates (mGy h<sup>-1</sup>) |  |  |  |  |
| --- | --- | --- | --- | --- | --- | --- | --- |
|  |  |  | 1 | 10 | 20 | 40 | 100 |
| ATP-dependent DNA helicase DDM1 | *DDM1* | MA_42406g0010 | -0.54 | -0.50 | -1.04 | -1.19 | -2.02 |
| DNA (cytosine-5)-methyltransferase CMT2 | *CMT2* | MA_173651g0020 | -0.59 | -0.69 | -1.08 | -1.42 | -2.34 |
| DNA (cytosine-5)-methyltransferase CMT3 | *CMT3* | MA_173651g0010 | -0.40 | -0.60 | -0.83 | -1.26 | -1.81 |
| Histone chaperone ASF1B | *ASF1B* | MA_73031g0010 | -0.39 | -0.64 | -0.60 | -1.48 | -1.94 |
| DNA (cytosine-5)-methyltransferase 1 | *DMT1* | MA_88843g0010 | -0.37 | -0.75 | -1.02 | -1.59 | -2.48 |
| Histone H2A | *HTA5* | MA_601125g0010 | -0.69 | -0.89 | -1.32 | -0.92 | -2.28 |
| HTB11 | *HTB11* | MA_454575g0010 | -0.74 | -0.99 | -1.26 | -1.92 | -2.01 |
| Histone-lysine N-methyltransferase, H3 lysine-9 specific SUVH5 | *SUVH5* | MA_790260g0010 | -0.79 | -0.88 | -1.56 | -2.25 | -2.80 |
| Histone-lysine N-methyltransferase SUVR4 | *SUVR4* | MA_106792g0010 | -0.26 | -0.53 | -0.84 | -1.51 | -1.79 |
| Histone-binding protein MSI1 | *MSI1* | MA_10432677g0010 | -0.22 | -0.69 | -0.86 | -1.55 | -2.17 |

### Altered expression of UPR- and vesicular transport-related genes in response to gamma irradiation

Genes in the UPR pathway showed significantly changed expression (Table 5, Table S8). The ER membrane protein gene *SRC2*, *YKT61,* a core component of the membrane fusion machinery SNARE complex, were upregulated at ≥ 40 mGy h^-1^ and the *Arabidopsis* type II Metacaspase homolog *AMC6* were upregulated at ≥ 20 mGy h^-1^. Additionally, small HSPs genes (*sHSP*s), such as *HSP17.6c* and *HSP21* were upregulated even at ≥ 10 mGy h^-1^ and ≥ 20 mGy h^-1^, respectively. On the other hand, *VACUOLAR PROTEIN SORTING 41* (*VPS41*) being essential for vacuole biogenesis and fusion, and several genes encoding heat shock proteins (*HSP*s) such as *HSP70* and *HSP90,* were downregulated at ≥ 40 mGy h^-1^. Moreover, the E3 ubiquitin-protein ligase UPL2 and RBR-type E3 ubiquitin transferase transcripts were downregulated along with many other proteasome components (Table 5, Table S8).

**Table 5.** Differentially expressed unfolded protein response-(UPR) genes in clonal stem cells of Norway spruce after exposure to 144 h of gamma radiation at different dose rates. Red and blue highlights denote upregulation and downregulation, respectively, compared to unexposed control (0 mGy h^-1^). Green borders denote significant fold changes (p□≤□0.05).

| Name | Name abbreviation | Gene Id | Fold change (Log<sub>2</sub>) in dose rates (mGy h<sup>-1</sup>) |  |  |  |  |
| --- | --- | --- | --- | --- | --- | --- | --- |
|  |  |  | 1 | 10 | 20 | 40 | 100 |
| E3 ubiquitin-protein ligase UPL2 | *UPL2* | MA_10436069g0010 | -0.45 | -0.43 | -1.32 | -0.86 | -1.52 |
| RBR-type E3 ubiquitin transferase |  | MA_10053g0010 | -0.47 | -0.26 | -0.96 | -1.06 | -1.82 |
| MCP2c | *AMC6* | MA_10430487g0010 | 0.10 | 0.48 | 1.13 | 1.81 | 2.31 |
| Hsp70-Hsp90 organizing protein 3 | *HOP3* | MA_867694g0010 | -0.10 | -0.33 | -0.31 | -0.47 | -0.92 |
| 17.6 kDa class I heat shock protein 3 | *HSP17.6C* | MA_10427493g0010 | 0.66 | 1.22 | 1.00 | 2.77 | 3.30 |
| HSP21 | *HSP21* | MA_10434235g0010 | 0.22 | 0.55 | 1.20 | 1.74 | 1.87 |
| Heat shock 70 kDa protein 10, mitochondrial | *HSP70-10* | MA_79699g0010 | -0.23 | -0.43 | -0.26 | -1.06 | -1.60 |
| Heat shock protein 90-1 | *HSP90-1* | MA_10861g0010 | 0.24 | -0.41 | -0.30 | -0.80 | -1.60 |
| HSP90.5 | *HSP90-5* | MA_10431770g0010 | 0.06 | -0.24 | -0.34 | -0.89 | -1.34 |
| Heat shock protein 90-6, mitochondrial | *HSP90-6* | MA_82160g0010 | -0.14 | -0.24 | -0.27 | -0.77 | -1.25 |
| Protein SRC2 homolog | *SRC2* | MA_5321101g0010 | -0.16 | -0.11 | 0.52 | 0.99 | 1.52 |
| SNARE-like superfamily protein | *YKT61* | MA_252882g0010 | -0.01 | 0.20 | 0.46 | 0.83 | 1.01 |
| Protein SUPPRESSOR OF K(+) TRANSPORT GROWTH DEFECT 1 | *SKD1* | MA_465725g0010 | 0.02 | 0.20 | 0.33 | 0.68 | 0.78 |
| Vacuolar protein sorting-associated protein 41 homolog | *VPS41* | MA_10436506g0010 | -0.28 | -0.22 | -0.43 | -0.53 | -0.93 |

### Gamma radiation-induced altered expression of cell wall- and hormone-related genes

Many cell wall-related genes were significantly regulated in response to the gamma irradaiton (Table 6, Table S9). Pectinacetylesterase, pectinmethylestarase, pectinlyase and pectinestarase-encoding genes related to cell wall modifications, as well as the xyloglucan galactosyltransferase *MUR3* and the cellulose synthase gene *CESA8,* were mostly upregulated at 40-100 mGy h^-1^. Cellulose synthase gene *CSLE1* was upregulated at 100 mGy h^-1^ (Table 6, Table S9).

**Table 6.** Differentially expressed cell wall related genes in clonal stem cells of Norway spruce after exposure to 144 h of gamma radiation at different dose rates. Green borders denote significant fold changes (p□≤□0.05). Green borders denote significant fold changes (p□≤□0.05).

| Name | Name abbreviation | Gene Id | Fold change (Log<sub>2</sub>) in dose rates (mGy h<sup>-1</sup>) |  |  |  |  |
| --- | --- | --- | --- | --- | --- | --- | --- |
|  |  |  | 1 | 10 | 20 | 40 | 100 |
| Pectin lyase-like superfamily protein |  | MA_98717g0010 | 0.19 | 0.84 | 1.13 | 1.87 | 2.13 |
| Pectin lyase-like superfamily protein |  | MA_9165956g0010 | 0.55 | 1.46 | 1.02 | 1.65 | 3.20 |
| Probable pectinesterase 53 | *PME53* | MA_10428207g0020 | 0.72 | 1.07 | 0.58 | 2.01 | 2.90 |
| Pectinesterase 2 | *PME2* | MA_84680g0010 | 0.87 | 1.09 | 0.63 | 1.23 | 2.44 |
| Pectin acetylerase 11 | *PAE11* | MA_10376612g0010 | -0.13 | 0.50 | 0.55 | 1.66 | 1.69 |
| Pectinesterase inhibitor 9 | *PMEI9* | MA_29344g0010 | -0.24 | 0.20 | 0.58 | 0.64 | 0.95 |
| pectin methylesterase PCR fragment F | *ATPMEPCRF* | MA_10332149g0010 | 0.24 | 0.45 | 0.12 | 2.60 | 2.36 |
| Pectinesterase |  | MA_67675g0010 | -0.08 | -0.24 | -0.21 | -0.72 | -1.24 |
| Pectinesterase 11 | *PE11* | MA_7274893g0010 | 0.41 | 0.40 | 0.76 | 0.97 | 1.32 |
| pectin methylesterase PCR fragment F | *ATPMEPCRF* | MA_177599g0010 | 0.05 | -0.48 | -0.70 | -0.16 | -1.23 |
| Probable pectinesterase 53 | *PME53* | MA_99816g0010 | 0.15 | 0.57 | 1.07 | 1.71 | 2.20 |
| Pectinesterase 11 | *PE11* | MA_87592g0010 | 0.62 | 1.10 | 1.02 | 2.33 | 2.89 |
| Pectinesterase 11 | *PE11* | MA_304854g0010 | 0.49 | 1.20 | 0.91 | 2.31 | 3.15 |
| Pectinesterase 11 | *PE11* | MA_18312g0010 | 0.67 | 1.14 | 0.39 | 1.39 | 2.11 |
| pectin methylesterase PCR fragment F | *ATPMEPCRF* | MA_158002g0020 | -0.28 | 0.24 | 0.03 | 1.90 | 1.29 |
| Xyloglucan galactosyltransferase MUR3 | *MUR3* | MA_13968g0020 | 0.11 | 0.45 | 0.62 | 1.21 | 1.28 |
| Xyloglucan galactosyltransferase MUR3 | *MUR3* | MA_100374g0010 | 0.42 | 0.37 | 0.43 | 1.20 | 1.83 |
| Xyloglucan galactosyltransferase MUR3 | *MUR3* | MA_7543346g0010 | -0.11 | 0.42 | -0.05 | 1.54 | 1.66 |
| Cellulose synthase-like protein E1 | *CSLE1* | MA_195271g0010 | 0.81 | 1.42 | 1.06 | 0.46 | 2.13 |
| Cellulose synthase | *CESA1* | MA_10430668g0010 | -0.09 | -0.45 | -0.48 | -0.94 | -1.17 |
| Cellulose synthase | *CESA8* | MA_8216532g0010 | 0.69 | 1.12 | 0.87 | 0.08 | 1.40 |

Also, significant enrichment of GO terms related to hormone biosynthesis and signaling were observed (Table 7, Table S10). Most DEGs in jasmonic acid biosynthetic process, response to jasmonic acid, salicylic acid-mediated signaling pathway and auxin efflux showed significant regulation from 40 mGy h^-1^, but a few genes even at ≥ 10 mGy h^-1^ (Table 7, Table S10). DEGs encoding auxin-responsive factors (ARFs), such as *ARF16*, *ARF6* and *JAR1*, an auxin-responsive GH3 family protein, and the early auxin-responsive protein *SMALL AUXIN UP RNA 32* (*SAUR32*) were upregulated at ≥ 10 mGy h^-1^ whereas other auxin response and transport genes, *IAA13*, *IAA26*, *PIN2* and *PIN3* were downregulated at ≥ 40 mGy h^-1^ (Table 7, Table S10). Genes encoding ethylene-responsive transcription factors, such as the *ERF53, ERF1A*, *ERF4*, and *RELATED TO AP2-11* (*RAP2-11*), showed upregulation at ≥ 40 mGy h^-1^, whereas *ERF013* showed upregulation at ≥ 20 mGy h^-1^. The cytokinin-biosynthesis gene *CYTOKININ HYDROXYLASE*, the cytokinin degradation gene *CYTOKININ DEHYDROGENASE* 5 (*CKX5*), and the gibberellin (GA) biosynthesis genes *ENT-KARURENE SYNTHASE* (*GA2*) and *GIBBERELLIN 20-OXIDASE 2* (*GA20OX2*) were all downregulated at ≥ 40 mGy h^-1^.

**Table 7.** Differentially expressed hormone biosynthesis and signalling genes in clonal stem cells of Norway spruce after exposure to 144 h of gamma radiation at different dose rates. Red and blue highlights denote upregulation and downregulation, respectively, compared to unexposed control (0 mGy h-1). Green borders denote significant fold changes (p□≤□0.05).

| Name | Name abbreviation | Gene Id | Fold change (Log<sub>2</sub>) in dose rates (mGy h<sup>-1</sup>) |  |  |  |  |
| --- | --- | --- | --- | --- | --- | --- | --- |
|  |  |  | 1 | 10 | 20 | 40 | 100 |
| Auxin response factor 16 | *ARF16* | MA_2421g0010 | -0.07 | 0.51 | 0.74 | 1.55 | 1.98 |
| Auxin response factor 6 | *ARF6* | MA_73460g0010 | 0.06 | 0.17 | 0.34 | 0.63 | 0.94 |
| Auxin-responsive protein IAA13 | *IAA13* | MA_84793g0010 | 0.41 | -0.49 | -0.46 | -1.27 | -2.21 |
| Auxin-responsive protein IAA26 | *IAA26* | MA_123046g0010 | -0.04 | -0.66 | -1.10 | -1.83 | -2.34 |
| Auxin-responsive protein IAA33 | *IAA33* | MA_9357208g0010 | 0.33 | 0.63 | 1.07 | 1.98 | 2.14 |
| WAV6 | *PIN2* | MA_61553g0020 | 0.23 | -0.40 | -0.22 | -0.73 | -1.11 |
| Auxin efflux carrier component 3 | *PIN3* | MA_61553g0010 | 0.01 | -0.56 | -0.18 | -0.83 | -0.86 |
| Auxin-responsive protein SAUR32 | *SAUR32* | MA_5864g0010 | 0.77 | 1.39 | 1.11 | 1.53 | 3.17 |
| Auxin-responsive GH3 family protein | *JAR1* | MA_376158g0010 | -0.34 | 1.35 | 1.18 | 1.48 | 1.82 |
| Cytokinin dehydrogenase 5 | *CKX5* | MA_10433485g0010 | -0.54 | -1.11 | -1.96 | -1.96 | -2.95 |
| Cytokinin hydroxylase | *CYP735A2* | MA_9176g0010 | 0.11 | -0.66 | 0.32 | -0.88 | -1.76 |
| Ethylene-responsive transcription factor ERF013 | *ERF013* | MA_3758g0010 | 0.39 | 0.46 | 0.74 | 1.46 | 2.04 |
| Ethylene-responsive transcription factor ERF053 | *ERF053* | MA_2446g0010 | 0.24 | 0.32 | 0.81 | 0.94 | 1.89 |
| Ethylene-responsive transcription factor 1A | *ERF1A* | MA_27309g0010 | 0.37 | 0.80 | 0.26 | 1.05 | 2.00 |
| Ethylene-responsive transcription factor 4 | *ERF4* | MA_81029g0010 | -0.11 | 0.12 | 0.11 | 0.89 | 1.14 |
| Ethylene-responsive transcription factor RAP2-11 | *RAP2-11* | MA_65190g0010 | 0.07 | 0.55 | 0.82 | 1.70 | 1.90 |
| Ethylene-responsive transcription factor ESR2 | *ESR2* | MA_77420g0010 | -0.69 | -0.79 | -1.28 | -1.77 | -2.29 |
| Ethylene-responsive transcription factor ERF055 | *ERF055* | MA_2040g0010 | -1.07 | -0.94 | -1.39 | -1.46 | -1.62 |
| KS1 | *GA2* | MA_57103g0010 | -0.06 | -0.83 | -1.25 | -2.27 | -2.82 |
| Gibberellin 20 oxidase 2 | *GA20OX2* | MA_281894g0010 | 0.44 | 0.03 | -0.52 | -1.60 | -2.42 |

### Post-irradiation expression of selected DNA repair and UPR-related genes

To assess how gamma radiation-regulated DDR-, and UPR-related genes were affected post-irradiation, the expression of selected genes at the end of the irradiation (T1) and in a subsequent proliferation (P2) stage were analysed by qPCR (Fig. 4). Whereas *RAD5* and *SOG1* showed a gamma dose rate-dependent expression at T1, with significant downregulation at 40-100 mGy h^-1^, no significant difference from the unexposed control was observed in P2. Expression of *RAD50, GR1* and *XRCC3* showed much variation at both stages with no clear dose rate-dependent effect. UPR-related genes exhibited significant dose rate-dependent effects mostly at T1 but generally not in P2. Of these genes, *HSP70* and *HSP90* were significantly downregulated at 40-100 mGy h ^-1^ at T1 and *HSP90* was significantly upregulated at all dose rates in P2, up to 4-fold compared to T1. *RBRE3* was significantly downregulated at ≥ 20 mGy h^-1^ at T1 but no dose-dependent effect was observed in P2.

**Fig. 4.**
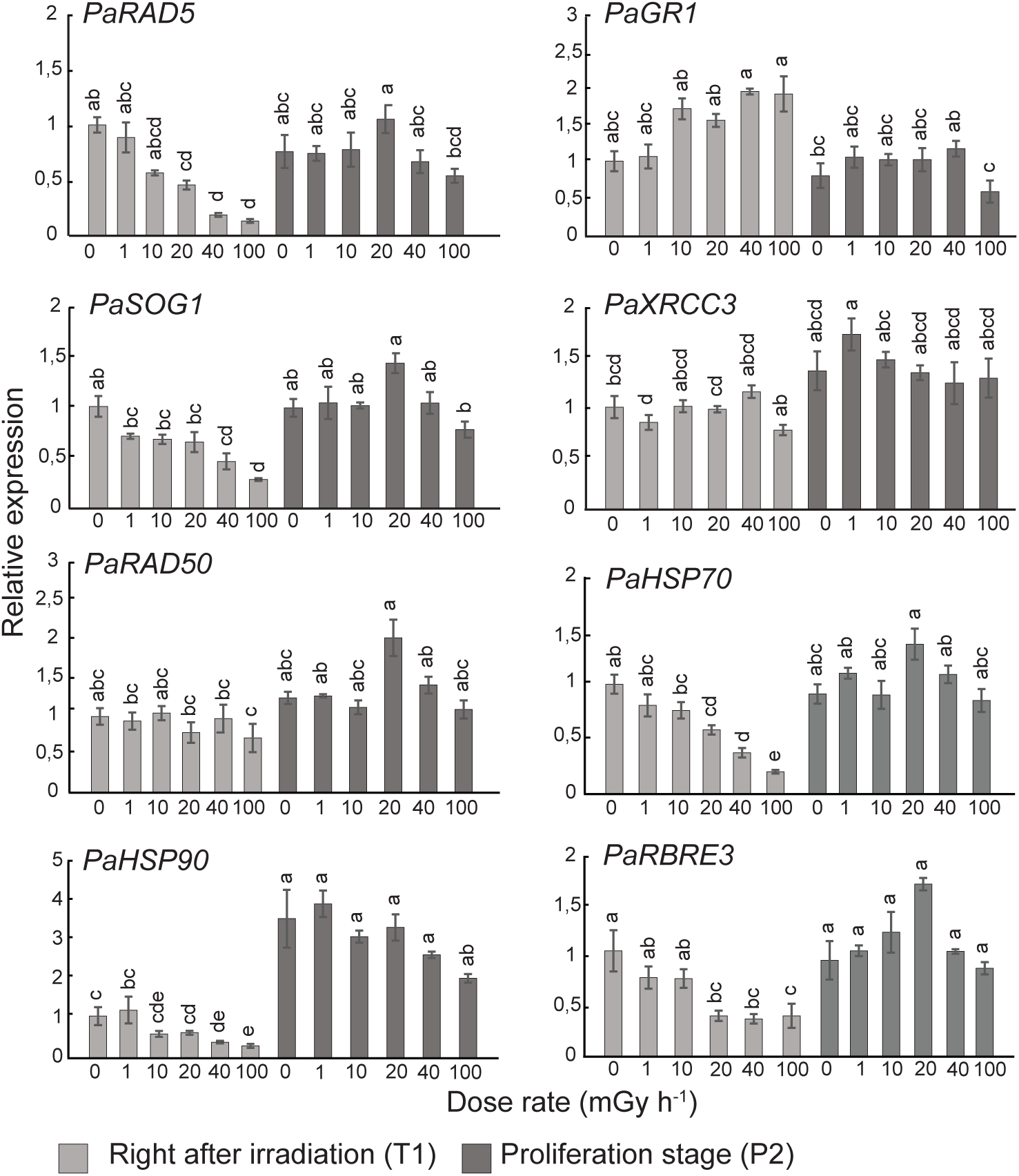
Relative transcript levels of DNA damage repair- (*RAD5*, *GR1*, *SOG1*, *XRCC3*, *RAD50)* and unfolded protein response (UPR; *HSP70*, *HSP90*, *RBRE3*)-related genes after 144 h of gamma irradiation of clonal stem cells of Norway spruce and at 7 weeks post-irradiation (proliferation stage 2). Transcript levels were normalised against *ACTIN*, *TUBULIN*, and *ELONGATION FACTOR 1* and shown relative to the unexposed controls. The results are mean ± SE (n = 4 with 3-4 technical replicates). Different letters within each gene indicate significant difference (p ≤ 0.05) based on two-way analysis of variance using log-transformed data, followed by Tukey’s post-hoc test.

## Discussion

Adverse effects of ionising radiation on plant growth and development including damage to the SAM-harbouring shoot tips in radiation-vulnerable conifers are well known (Caplin and Wiley, 2018 and references therein; (Blagojevic et al. 2019a). Despite the significance of the stem cells of the SAMs for growth, reproduction and survival of plants, detailed molecular effects of ionising radiation on such stem cells are virtually unexplored. Here we employed *in vitro*-grown clonal stem cells of Norway spruce to assess the effect of gamma radiation on their proliferation and stemness as well to characterise such cells as a model to investigate molecular toxicity of gamma radiation in ecologically and economically important conifers.

Although no visible damage or reduced cell proliferation of the stem cells were observed after 144 h of gamma irradiation or during the post-irradiation period, damage to DNA, nuclei and mitochondria as well as accumulation of amorphous materials in cells at all dose rates (Fig. 1, 2, Fig S1) demonstrate profound adverse cellular effects. Differences in abundance of specific cell wall components between gamma-irradiated and control cells also suggest ionising radiation-induced cell wall modifications (Fig. S2). The lack of detection of methyl-esterified HG pectin at the end of the gamma irradiation and higher abundance of un-methyl-esterified HG post-irradiation compared to in unexposed control cells are consistent with the previously reported stress-induced de-methyl-esterification (Lee et al. 2017; Novaković et al. 2018; Rui and Dinneny 2020). Also, detection of xyloglucan and arabinogalactan protein in gamma-irradiated cells only is in line with previous reported changes in the hemicellulose fraction and induction of arabinogalactan protein in response to abiotic stress (Le Gall et al. 2015). Yet, in spite of the observed adverse cellular effects, a fraction of the stem cells obviously survived and retained their stemness and accordingly continued to proliferate (Fig. 1a, Fig S1), suggesting mobilisation of repair and protection systems at least to some extent. This at least partly resembles the situation with survival and activation of axillary meristems despite mortality of the SAM (manifested as lack of apical dominance) in conifers in radionuclide-contaminated areas (Caplin and Wiley 2018; Nybakken et al. 2023).

In addition to direct ionising of vital molecules, gamma radiation results in ROS accumulation, which can cause persistent damage to macromolecules but also induce protective antioxidant biosynthesis (You and Chan 2015); (Sauer, Wartenberg, and Hescheler 2001). The massive upregulation of antioxidant genes encoding peroxidases, glutathione-s-transferases and lignans (Table 2. Table S4) is consistent with this and may at least partly explain the survival of at least a fraction of the stem cells. However, the activity of antioxidants was obviously not sufficient to neutralise the oxidative stress, as judged from the gamma radiation-induced dose rate-dependent DNA and organelle damage. The downregulation of genes encoding SOD (Table 2, Table S4), likely contributed to the observed organelle damage. These effects of the 144-h of gamma irradiation on the stem cells of Norway spruce are comparable to observed effects of gamma irradiation of the same duration in Norway spruce and Scots pine seedlings (Blagojevic et al. 2019a) and similar to reported effects of high levels of ionising radiation on other plant species such as *A. thaliana* (Gill and Tuteja 2010; Gill et al. 2015).

Extensive DNA damage is supposed to activate the DDR pathway (Kim et al. 2019). However, in this study, we found overall transcriptional repression of DDR genes, including crucial components of nucleotide excision repair (NER), homologous recombination (HR), replication repair and activation of cell cycle checkpoints (Table 3, Tables S5, S6), suggesting an overall inhibition of the DNA repair consistent with the observed DNA damage in the stem cells (Fig. 1b). In *A. thaliana* plants, cell cycle arrest through a SOG1- and ATM/ATR-dependent DDR pathway is a well-known indicator of DNA damage (Kim et al. 2019; Yoshiyama et al. 2014). Hence, the transcriptional repression of *SOG1* from the lowest dose rates and dose rate-dependent downregulation of cyclins and CDKs (Table 3, Table S5) in the stem cells in our study, imply a negative effect on the cell cycle, with disruption of cell division and DNA replication (Adachi et al. 2011).

The reduced expression of cell division-related genes is consistent with the downregulation of genes related to biosynthesis of cytokinin and gibberellin and a range of auxin-responsive and auxin transport genes, as well as the upregulation of genes encoding stress-related ethylene-responsive transcription factors (Table 7, Table S10). However, although biosynthesis of the main cell division-stimulating hormone cytokinin might be compromised in the irradiated proliferating cells, the downregulation of the cytokinin-degradation gene *CKX* and upregulation of the cytokinin-activation gene *LOG*, may suggest that stability of active cytokinin was maintained (Table S10).

Exposure to ionising radiation can cause genomic instability and long-term effects that can be associated with epigenetic mechanisms such as DNA methylation and modifications of histone tails (Merrifield and Kovalchuk 2013; Braszewska-Zalewska et al. 2014). Multifold repression of multiple genes involved in DNA methylation, histone modification, and nucleosome assembly in the gamma-irradiated stem cells (Table 4, Table S7) suggest altered DNA methylation status and modified chromatin dynamics leading to stress-induced genomic instability. Downregulation of *DDM1*, which is involved in DNA repair and methylation, in the stem cells in our study resembled the situation with hypomethylation of DNA and accumulation of DNA damage in UV-B-exposed *A. thaliana* (Müller-Xing et al. 2014). Additionally, the suppression of other genes encoding integral components of the global DNA methylation pathway such as DMT1, CMT2 and 3, possibly indicates altered methylation status in the gamma-exposed stem cells compared to the unexposed controls (Feng et al., 2010; Shen et al. 2014). Moreover, since the histone proteins are the building blocks of the nucleosomes, whose synthesis, modification and exchange are required to maintain appropriate transcriptional properties (Zhou et al. 2015), the repression of *HTA5* and *HTB11* expression by several folds in the stem cells indicates that gamma irradiation has a negative effect on the overall synthesis of H2A and H2B (Zhou, et al. 2015; Braszewska-Zalewska et al. 2014); (Moreno-Romero et al. 2012). This could mean a reduced availability of core histone components affecting chromatin dynamics and transcriptional regulation of genes. However, as H2A modification is also associated with DDR, this downregulation may also have contributed to the ineffective DDR in the gamma-irradiated stem cells.

Upon stress exposure, UPR is induced to cope with misfolded and unfolded proteins in ER by activating molecular chaperons and degradation of misfolded proteins to restore the homeostasis (Howell 2013). Activation of UPR-transducers such as bZIP17/28/60, NAC017/062/089/103 and others are crucial in this process (Nawkar et al. 2018). In the gamma-irradiated stem cells, significant upregulation of *bZIP17* (Table 5, Table S8) at all dose rates is a direct indication of UPR being a fundamental response in ionising radiation-induced stress. Additionally, genes encoding chaperons such as HSPs in ER and the interaction between HSP90 and the 26s proteasome machinery are crucial for the assembly and maintenance of the proteasome required for stress-mediated protein folding (Nishizawa-Yokoi et al. 2010; Kurepa et al. 2009). Hence, downregulation of multiple homologs of *HSP*s, *HOP*s and *E3 LIGASES* from 20 mGy h^-1^ (Table 5) indicated severely compromised protein folding machinery (Sharma et al. 2016; Peralta et al. 2013; Mazzucotelli et al. 2006). Some HSPs also function in coordination with other molecular chaperones and protect misfolded proteins targeted to ATP-dependent refolding or degradation (Bernfur et al. 2017), and are known to provide stress tolerance in *A. thaliana* (Sun et al. 2002). Thus, significant induction of *HSP17.8* and *HSP21* expression in response to the gamma radiation (Table 5, Table S8) indicated that components of the stress response mechanism involved in the maintenance of the structural integrity of proteins are to some extent in action to provide some protection against the gamma radiation-related damage.

Continued stem cell proliferation post-irradiation was consistent with the altered post-irradiation expression (stage P2) of specific DDR- and UPR-related genes as revealed by RT-qPCR analyses compared to at the end of the gamma-irradiation (stage T1). Whereas the DDR-related genes *RAD51* and *SOG1* and the UPR-related genes *HSP70* and *HSP90* showed downregulation at the T1 stage, no significant regulation of *RAD51* and *SOG1* and upregulation of *HSP90* at the P2 stage imply that the stem cells were able to recover from the radiation-induced transcriptional repression post-irradiation. Moreover, despite the substantial cellular and DNA damage, after the transfer of stem cells to maturation media for embryogenesis, surviving cells were even able to develop somatic embryos at all gamma dose rates during the post-irradiation period (Fig. S1). Although such somatic embryo cultures may commonly generate a proportion of abnormal embryos (Alvarez et al. 2015), the lack of normal cotyledon number (7-8) at all gamma dose rates in contrast to the control cultures, indicates some adverse after-effect of the gamma radiation. Nevertheless, the post-irradiation embryo development at all dose rates could be due to that the surviving cells then had mobilised sufficient repair and protection systems enabling them to complete the embryogenesis on induction media albeit with some embryo abnormalities.

## Conclusions

Collectively, the results demonstrate that at least a fraction of the pluripotent stem cells could retain their stemness after exposure to low to moderate levels (1-100 mGy h^-1^ for 144 h) of gamma radiation despite substantial DNA and organelle damage. The RNA-seq analyses revealed massive gamma radiation-induced changes in gene expression towards stress management with upregulation of specific antioxidant genes, protein transport and degradation, but also downregulation of DDR, DNA replication, cell division and protein-folding (UPR) related genes. Yet, the normalisation of the expression of the DDR-related genes *RAD5*, *SOG1* and the UPR-related genes *HSP90*, *HSP70* and *RBRE3* during the post-irradiation proliferation together with the observed continued cell proliferation and somatic embryo development after transfer to maturation media, demonstrate that the stem cells are able to recover from the gamma radiation-induced stress. Thus, although they sustained comprehensive damages to an extent that a portion of them died, the surviving cells maintained their stemness and some were capable of embryo formation.

## CRediT authorship contribution statement

**Payel Bhattacharjee**: Writing - original draft, Writing - review and editing, Conceptualisation, Investigation, Formal analysis, Data curation, Validation, Visualisation. **YeonKyeong Lee**: Writing - original draft, Writing - review and editing, Conceptualisation, Investigation, Formal analysis, Data curation, Validation, Visualisation. **Marcos Viejo**: Writing - review and editing, Investigation. **Gareth B Gillard**: Writing - review and editing, Data curation, Formal analysis. **Simen Rød Sandve**: Writing - review and editing, Supervision. **Torgeir R Hvidsten**: Writing - review and editing, Supervision. **Brit Salbu**: Writing - review and editing, Conceptualisation, Funding acquisition, Project administration **Dag A Brede**: Writing - review and editing, Conceptualisation, Investigation, Supervision. **Jorunn E. Olsen**: Writing - original draft, Writing - review and editing, Conceptualisation, Investigation, Validation, Visualisation, Funding acquisition, Project administration, Supervision.

## Data availability

The raw RNA sequence data is uploaded in ArrayExpress (accession number will be updated). Other data that support the findings of this study are available within the article, in the Supporting Information of this article and in online repositories. The names of the online repository and the link of the identifiers will be available in the Norwegian University of Life Sciences archive once the submission is approved.

## Acknowledgements

Sincere thanks to Tone I. Melby for skilful technical assistance with the RT-qPCR analyses.

