## Supplementary figures and images for "Gamma radiation-induced molecular toxicity and effects on pluripotent stem cells of the radiosensitive conifer Norway spruce (*Picea abies*)"

### Fig. S1

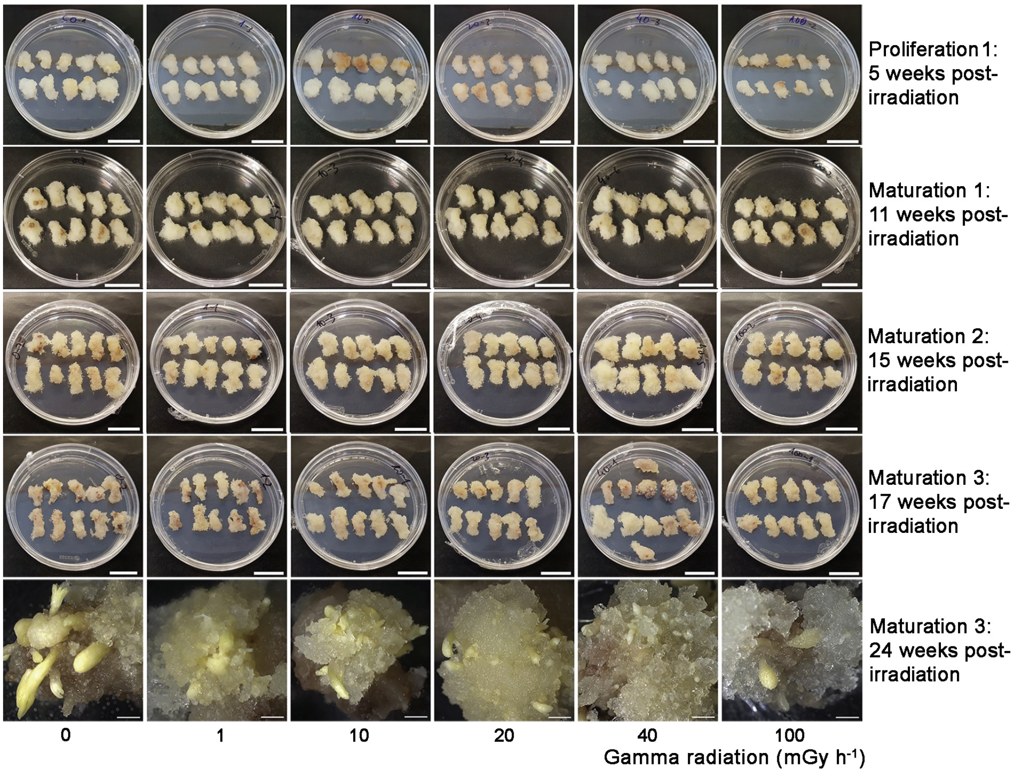


*Figure S1. Bhattacharjee and Lee et al.,2025.*

### Fig. S2

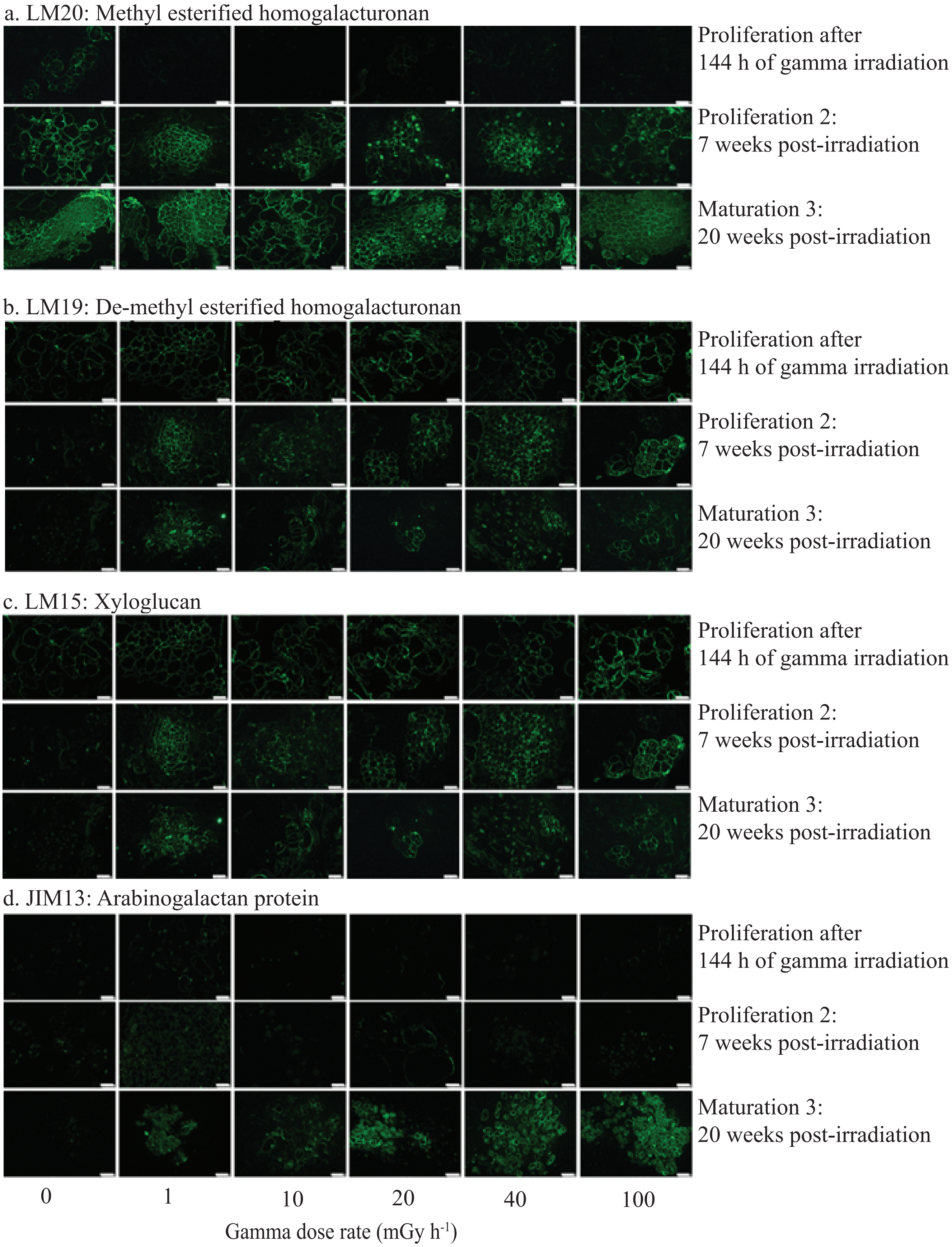


*Figure S2. Bhattacharjee and Lee et al.,2025.*

### Fig. S3

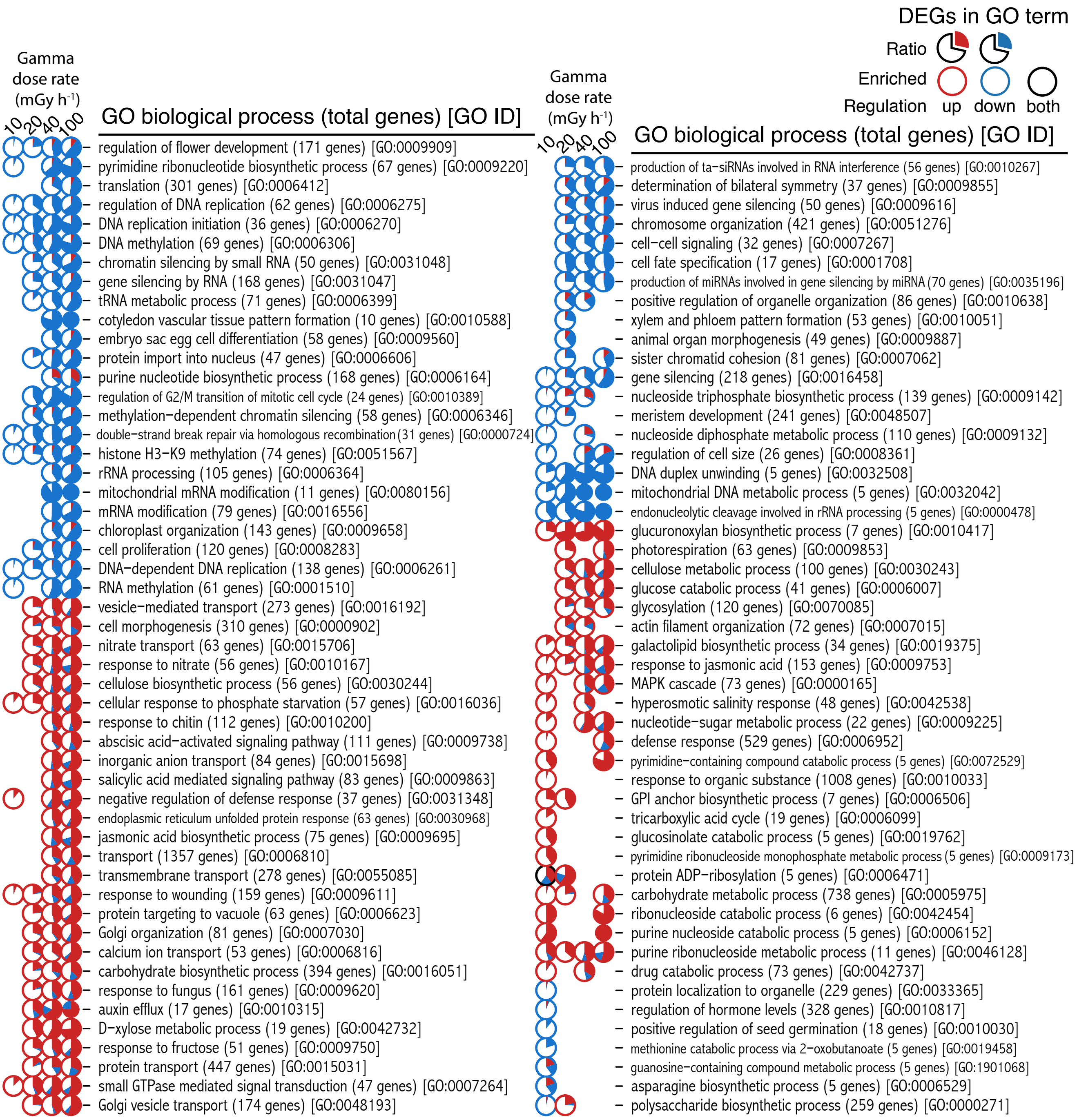


*Figure S3. Bhattacherjee, Lee et al., 2025*

### Fig. S4

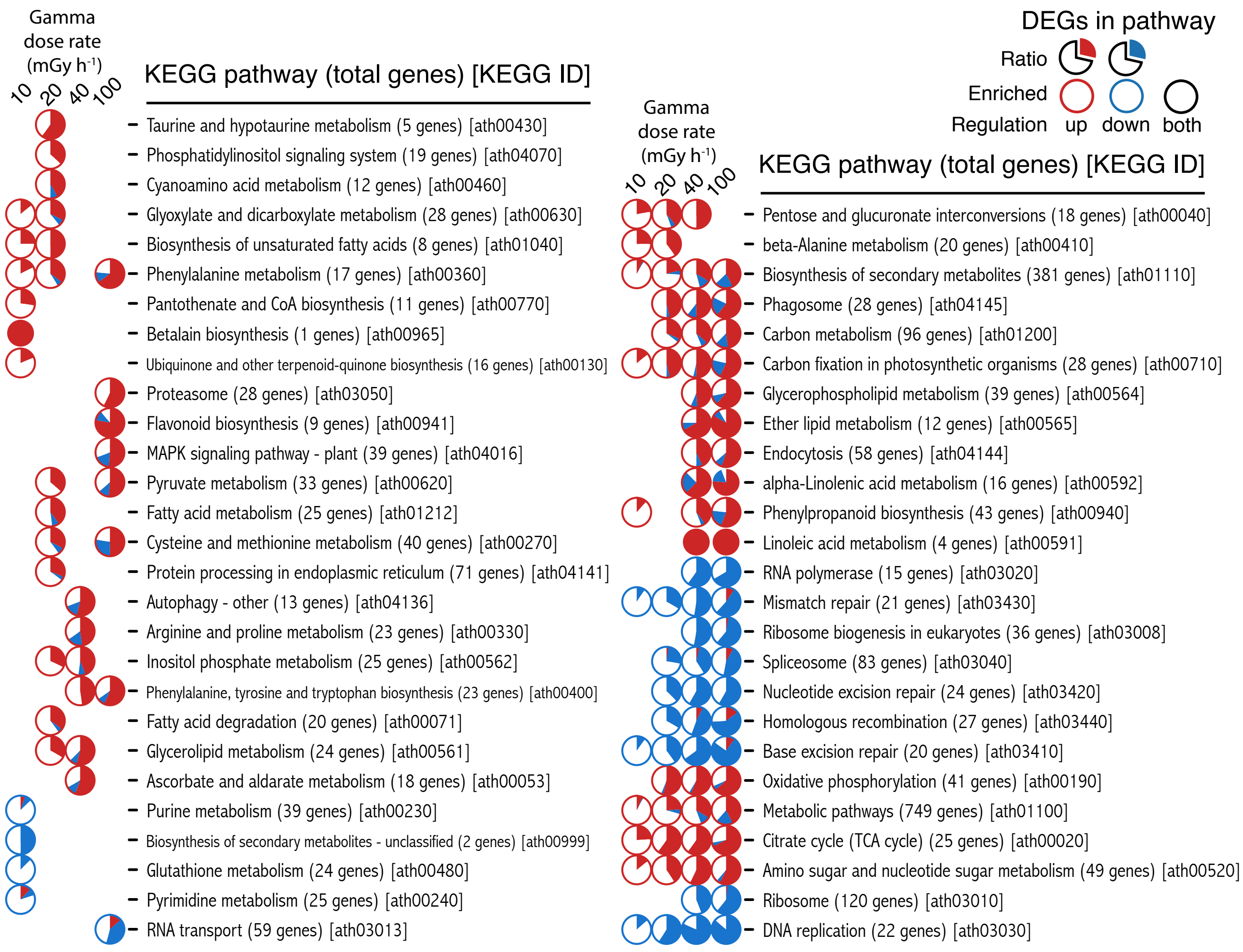


*Figure S4. Bhattacherjee, Lee et al., 2025*
