## Supplementary material for "Gamma radiation-induced molecular toxicity and effects on pluripotent stem cells of the radiosensitive conifer Norway spruce (*Picea abies*)": Figure legends

**Table headings for supplementary Tables**

**Table S1.** Cell wall polysaccharide-directed monoclonal antibodies (MAb) used to study effect of 144-h of gamma irradiation on cell wall composition in stem cells of Norway spruce.

**Table S2.** Primers (F: forward, R: reverse) for qPCR analysis of DNA repair- and unfolded protein response (UPR)-related genes after 144-h of gamma irradiation of stem cells of Norway spruce.

**Table S3.** The total number of somatic embryos formed post-irradiation after transfer to embryo-induction media following 144-h of gamma irradiation of proliferating cells on proliferation media. The results are from two repeated experiments with totally 40-60 irradiated cell aggregates per gamma dose rate.

**Table S4-11** Differentially expressed genes in selected gene groups in clonal stem cells of Norway spruce after 144 h of gamma irradiation. 4) Antioxidants, 5) DNA-repair, 6) cell cycle, 7) chromatin re-modellers, 8) unfolded protein response (UPR), 9) cell wall, 10) hormone, 11) transcription factors.

**Figure legends for Supplementary figures**

**Fig. S1.** Post-irradiation development of clonal stem cell aggregates and somatic embryos of Norway spruce following 144 h of gamma irradiation. Scale bars: 2:cm (4 upper panels) and 2 mm (lowest panel). Representative cell aggregates out of totally 40-60 irradiated ones per dose rates are shown.

**Fig. S2.** Micrographs of indirect immunofluorescence detection of cell wall components in proliferating stem cell aggregates of Norway spruce at the end of 144 h of gamma irradiation and during post-irradiation stages. Longitudinal 1 µm thick sections were probed with the monoclonal antibodies. Sections were made from 3 plants per gamma radiation dose rate. Scale bars: 50 µm.

**Fig. S3.** Gene Ontology (GO) enrichment analysis showing significant effects of 144-h irradiation with different gamma dose rates on expression of genes associated with various biological processes in genetically identical stem cells of Norway spruce, relative to unexposed control cells. For each gamma dose rate four repeated samples were analysed in duplicate by RNA sequencing.

**Figure S4.** KEGG ortholog enrichment and analysis of pathways (biological processes) significantly affected in genetically identical stem cells of Norway spruce exposed to different dose rates of gamma radiation for 144 h, relative to unexposed control cells. For each gamma dose rate four repeated samples were analysed in duplicate by RNA sequencing.
