## Supplementary material for "Gamma radiation-induced molecular toxicity and effects on pluripotent stem cells of the radiosensitive conifer Norway spruce (*Picea abies*)": Table S1

**Table S1.** Cell wall polysaccharide-directed monoclonal antibodies (MAb) used to study effect of 144-h of gamma irradiation on cell wall composition in stem cells of Norway spruce.

| **MAb** | **Cell wall polymer/epitope recognised** | **Reference** |
| --- | --- | --- |
| LM19 | Unesterified homogalacturonan | (Verhertbruggen et al. 2008) |
| LM20 | Methyl esterified homogalacturonan | (Verhertbruggen, et al. 2008) |
| LM15 | XXXG motif of xyloglucan | (Marcus et al. 2008) |
| JIM13 | Arabinogalactan protein | (Yates and Knox 1994) |
