## Supplementary material for "Gamma radiation-induced molecular toxicity and effects on pluripotent stem cells of the radiosensitive conifer Norway spruce (*Picea abies*)": Table S2

**Table S2.** Primers (F: forward, R: reverse) for qPCR analysis of DNA repair- and unfolded protein response (UPR)-related genes after 144-h of gamma irradiation of stem cells of Norway spruce.

| Primer name | Sequence (5'-3') |
| --- | --- |
| DNA repair |  |
| *PaRAD5F* | GTGTGCCCAGTATCAATAGTTGGTC |
| *PaRAD5R* | CCTAGAGTGGCATATGTAGTAATCACTATG |
| *PaRAD50F* | GCGAGATAGATATATTCAAAGTGTGTTCGC |
| *PaRAD50R* | GCTTTTGATTTTCATGTCATTAGACTCCTTC |
| *PaXRCC3F* | CAGCTGCAATTGGGTCATTCTTC |
| *PaXRCC3R* | ATTTCCGATTGGCCTGAAGCCC |
| *PaGR1F* | GCTTCGGAGCTTGAGCGTGAAAG |
| *PaGR1R* | CCAGTGTCAGGTGCCTGATAATCG |
| *PaSOG1F* | CCACTACGTTCATGGAACCAAC |
| *PaSOG1R* | ATCCGGATCTGCAGCCTGTG |
| UPR |  |
| *PaHsp70F* | GAAATGGGAAAATGCAGCAAGAGCTTTC |
| *PaHsp70R* | CTGAATTTTCAATTACTTTTGGGTTCTTCCC |
| *PaHsp90F* | CAAGCACAACGACGATGAGCAATAC |
| *PaHsp90R* | CTCCTCCAAGTACTCCAGATGGTC |
| *PaRBRE3F* | CCAAGGAGTGTTGTAGCTTTGCATG |
| *PaRBRE3R* | GAATTCTGAGGTCCCTCCCCAAG |
