## Supplementary material for "Gamma radiation-induced molecular toxicity and effects on pluripotent stem cells of the radiosensitive conifer Norway spruce (*Picea abies*)": Table S3

**Table S3.** The total number of somatic embryos formed post-irradiation after transfer to embryo-induction media following 144-h of gamma irradiation of proliferating cells on proliferation media. The results are from two repeated experiments with totally 40-60 irradiated cell aggregates per gamma dose rate.

| Dose rate (mGy h^-1^) | Number of embryos |
| --- | --- |
| 0 | 34 |
| 1 | 7 |
| 10 | 23 |
| 20 | 14 |
| 40 | 4 |
| 100 | 34 |
